# Directed evolution reveals the mechanism of HitRS signal transduction in *Bacillus anthracis*

**DOI:** 10.1101/2020.07.03.185868

**Authors:** Hualiang Pi, Michelle L. Chu, Samuel J. Ivan, Casey J. Latario, Allen M. Toth, Sophia M. Carlin, Gideon H. Hillebrand, Hannah K. Lin, Jared D. Reppart, Devin L. Stauff, Eric P. Skaar

**Affiliations:** Department of Pathology, Microbiology, & Immunology, Vanderbilt University Medical Center, Nashville, Tennessee, USA; Vanderbilt Institute for Infection, Immunology, & Inflammation, Vanderbilt University Medical Center, Nashville, Tennessee, USA; Department of Biology, Grove City College, Grove City, PA

**Keywords:** cell envelope stress, HAMP domain, genetic selection, phosphorylation, TCS signaling

## Abstract

Bacterial two <u>c</u>omponent <u>s</u>ystems (TCSs) have been studied for decades; however, most work has focused on individual domains or proteins. Systematic characterization of an entire TCS could provide a mechanistic understanding of these important signal transduction systems. Here, genetic selections were employed to dissect the molecular basis of signal transduction by the HitRS system that has been implicated in detecting cell envelope stress in the pathogen *Bacillus anthracis*. Numerous point mutations were isolated within HitRS, 17 of which were in a 50-residue HAMP domain. Mutational analysis revealed the importance of hydrophobic interactions within the HAMP domain and highlighted its essentiality in TCS signaling. In addition, these data defined residues critical for activities intrinsic to HitRS, uncovered specific interactions among individual domains and between the two signaling proteins, and revealed that phosphotransfer is the rate-limiting step for signal transduction. This study establishes the use of unbiased genetic selections to study TCS signaling, provides a comprehensive mechanistic understanding of an entire TCS, and lays the foundation for development of novel antimicrobial therapeutics against this important infectious threat.

## Introduction

The incidence of antibiotic resistant infections is rising globally, leading the world into a “post antibiotic era” and thus, the need to develop novel therapeutic interventions against bacterial pathogens is urgent. Emerging evidence suggests that targeting bacterial systems involved in stress response is a potential avenue for developing antimicrobials (Lee *et al*, 2009; Poole, 2014; Tkachenko, 2018). Therefore, identification and characterization of stress detection and detoxification mechanisms in pathogenic bacteria is of critical importance. *Bacillus anthracis* is a Gram-positive, spore-forming, facultative aerobe, and the causative agent of anthrax. *B. anthracis* spores can survive extreme temperatures, harsh chemical assaults, and nutrient-poor environments for many years (Goel, 2015). This pathogen is one of the few infectious agents that have been proven effective as weapons of bioterror and it causes a variety of infectious syndromes including cutaneous, gastrointestinal, and inhalation anthrax. Inhalation anthrax occurs when *B. anthracis* spores enter a host through the respiratory system before disseminating to the lymph nodes and is the most deadly form of anthrax with a mortality rate approaching 90% (Kamal *et al*, 2011). To survive interactions with the host immune system during infection, *B. anthracis* has developed comprehensive systems for stress detection and detoxification (Shatalin *et al*, 2008). Therefore, this intracellular pathogen is also an excellent model to study microbial stress responses.

Bacterial transcriptional changes in response to stress can be modulated by signal transduction systems known as two-<u>c</u>omponent <u>s</u>ystems (TCSs). TCSs detect a wide range of signals and stressors including pH, temperature, nutrient, light, small molecules, envelope stress, osmotic pressure, and the redox state (Brunskill & Bayles, 1996; Fournier & Klier, 2004; Giraudo *et al*, 1997; Martin *et al*, 1999; Recsei *et al*, 1986; Tkachenko, 2018; Yarwood *et al*, 2001). TCSs enable cells to sense, respond, and adapt to changes in their environment and regulate a wide variety of processes including virulence, sporulation, antibiotic resistance, nutrient uptake, quorum sensing, and membrane integrity (Hoch, 2017; Mike *et al*, 2014; Stauff & Skaar, 2009; Tierney & Rather, 2019). A prototypical TCS consists of a membrane-bound sensor protein (<u>h</u>istidine <u>k</u>inase, HK) and cytoplasmic response regulator (RR) (Bhate *et al*, 2015; Jacob-Dubuisson *et al*, 2018; Stock *et al*, 2000). A classic HK possesses five domains: a N-terminal <u>T</u>rans-<u>M</u>embrane domain (TM), a sensor domain, a HAMP domain that is commonly found in <u>H</u>istidine kinase, <u>A</u>denylyl cyclases, <u>M</u>ethyl-accepting chemotaxis protein, and <u>P</u>hosphatase, a DHp domain (<u>D</u>imerization and <u>H</u>istidine <u>p</u>hosphorylation), and a CA domain (<u>C</u>atalytic and <u>A</u>TP-binding) (Figure 1A) (Bhate *et al*., 2015; Jacob-Dubuisson *et al*., 2018; Stock *et al*., 2000). The latter two domains constitute the kinase core domains that harbor a number of well-characterized and conserved motifs (Figure S1). The DHp domain can be further divided into four subgroups based on sequence identity: HisKA, HisKA_2, HisKA_3, and his_kinase domains and about 80% of HKs contain a HisKA domain (Zschiedrich *et al*, 2016). The CA domain belongs to the HATPase_c domain family and exhibits a relatively slow ATPase activity (Wang *et al*, 2013). A typical RR consists of two domains: a phosphorylation receiver domain and an output effector domain (Figure 1A and S2), with more than 60% of the latter being a DNA-binding domain (Bhate *et al*., 2015; Jacob-Dubuisson *et al*., 2018; Stock *et al*., 2000). In the presence of a specific stimulus, the HK detects the signal via the sensor domain, transmits the signal onto the DHp domain through the HAMP linker, phosphorylates its own conserved His located in the DHp domain, and then transfers the phosphoryl group onto a conserved Asp in the receiver domain of the cognate RR. In the case of the RR being a transcriptional regulator, this phosphorylation event activates the RR, induces homodimerization of the receiver domain, stimulates binding to the target promoters, regulates target gene expression, and modulates cellular physiology in response to environmental stimuli. TCSs are present in nearly all sequenced bacterial genomes as well as some fungal, archaeal, and plant species but are absent in animals and humans (Bhate *et al*., 2015; Jacob-Dubuisson *et al*., 2018; Stock *et al*., 2000), making them attractive targets for antimicrobial therapeutics.

**Figure 1.**
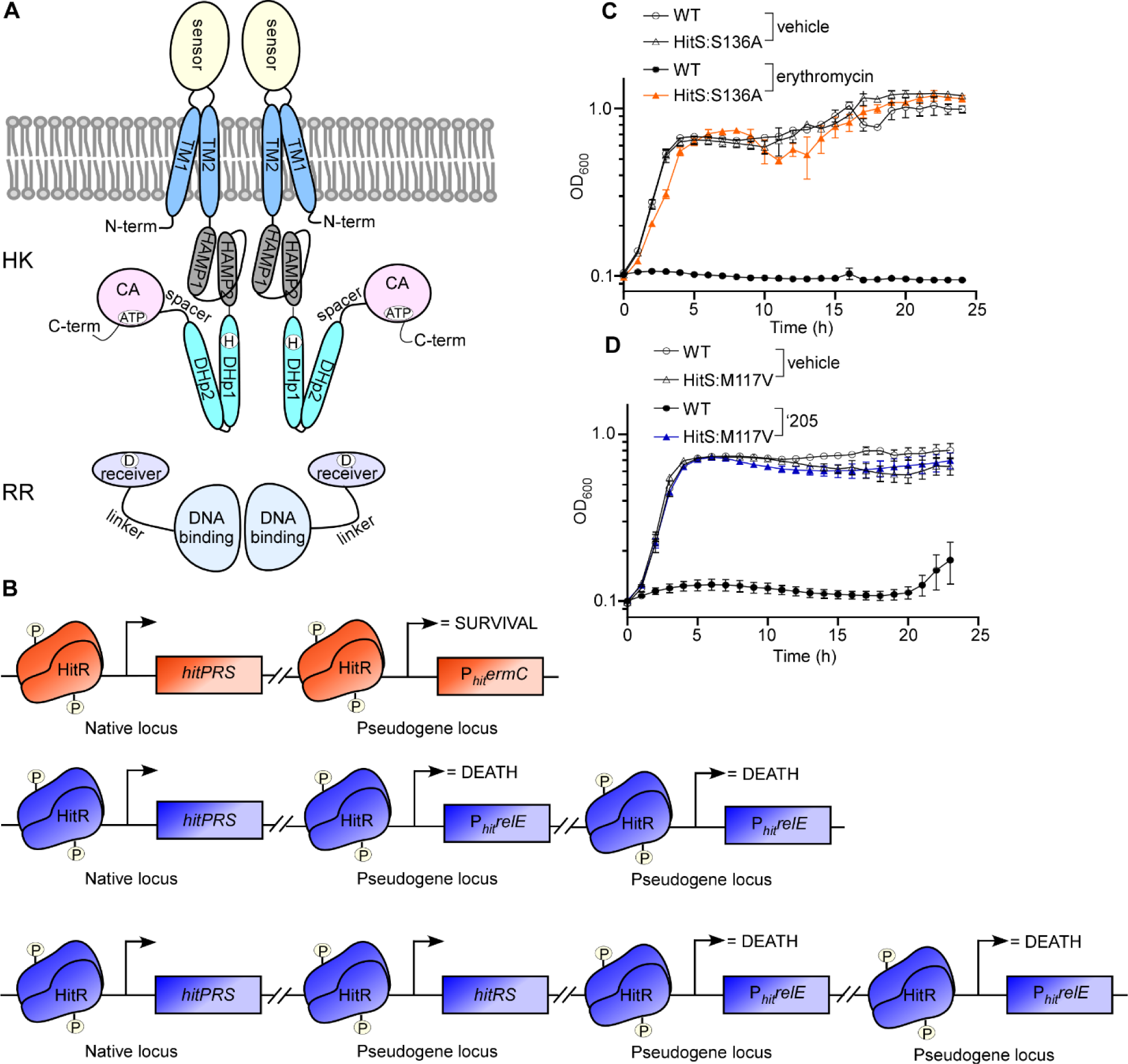
Genetic selection strategies to study HitRS signaling mechanism. (A) Schematics of the modular structure of a prototypical TCS. A classic histidine kinase (HK) consists of five domains: a N-terminal Trans-Membrane domain (TM), a sensor domain, a HAMP domain, a DHp domain, and a CA domain while a response regulator (RR) consists of two domains: a receiver domain and a DNA-binding domain. (B) Schematics for genetic selection strategies. To identify mutations that lead to constitutive activation of HitRS, an *ermC* strain shown in orange driven by a HitR-targeted promoter (P_*hit*_) was used. The two strains shown in blue (*relE* strains) were employed to isolate inactivating mutations within the TCS genes. (C) Growth kinetics of *ermC* expressing strains (WT and a representative activating mutant HitS:S136A) were monitored in the presence or absence of 20 µg ml^-1^ of erythromycin. (D) Growth kinetics of *relE* expressing strains (WT and a representative inactivating mutant HitS:M117V) were monitored in the presence or absence of 20 µM ‘205.

*B. anthracis* encodes approximately 45 TCSs, reflecting the complex environmental conditions encountered by this pathogen. A few *B. anthracis* TCSs have been studied (Laut *et al*, 2020; Mike *et al*., 2014; Stauff & Skaar, 2009), including a <u>h</u>eme <u>s</u>ensor <u>s</u>ystem (HssRS) that responds to changes in available heme and activates the expression of a heme efflux pump upon heme exposure, and a <u>H</u>ssRS interfacing <u>T</u>CS (HitRS) that senses cell envelope stress and activates an uncharacterized transporter system (HitP) (Mike *et al*., 2014; Stauff & Skaar, 2009). Although the nature of the activating signal of HitRS remains unclear, a high-throughput screen identified a series of cell-envelope acting compounds as inducers of HitRS (Mike *et al*., 2014; Mike *et al*, 2013), including the small synthetic compound VU0120205 (‘205) (Mike *et al*., 2013), nordihydroguaiaretic acid, which is an antioxidant that possesses activity against the cell membrane (Ooi *et al*, 2015), chlorpromazine, which is an antipsychotic drug that inhibits cell wall biogenesis (Klubes *et al*, 1971), targocil, which is an antibiotic that inhibits wall teichoic acid synthesis (Lee *et al*, 2010), and vancomycin, which inhibits Gram-positive cell wall biosynthesis and disrupts membrane integrity at low concentrations (Watanakunakorn, 1984). These compounds share little structural similarity but each is implicated in cell envelope stress suggesting that HitRS senses perturbations in the cell envelope.

In this study, genetic selection strategies were utilized to dissect the molecular mechanisms of signal transduction by HitRS. Numerous point mutations that lead to either inactivation or constitutive activation of the HitRS system were isolated. Representative mutations in each domain of these proteins were characterized biochemically to evaluate the effects of identified mutations on various activities required for signal transduction. These data uncovered the essential molecular determinants for HitRS signal sensing and promoter activation including: (i) four residues critical for the autokinase activity (S136/F149/V274/G309) besides the well conserved phosphoaccepting His and ATP-binding Asn, (ii) three residues essential for the phosphatase activity (S141/F149/R306), and (iii) five additional residues within HitR crucial for phosphotransfer and DNA-binding (F95/P106/P155/R192/Y222) besides the conserved phosphoaccepting Asp. In addition, our results revealed specific interactions among various domains, particularly the two kinase core domains and the two RR domains, and between HitR and HitS. Importantly, this study provides a detailed systematic characterization of TCS and expands our understanding of the molecular basis of signal transduction through TCS. Given that these signaling proteins are well conserved among distinct bacterial species, the described genetic selection and information obtained from it may be broadly applicable across multiple TCSs.

## Results

### Devising genetic selections to study the mechanism of HitRS signaling

To dissect the molecular determinants within HitRS that are required for signal sensing and promoter activation, two sets of genetic selections were performed. To isolate mutations that lead to constitutive activation of HitRS, we created a *B. anthracis* strain harboring the erythromycin resistance gene *ermC* driven by the HitRS promoter (P_*hit*_*ermC*) (Figure 1B). This strain was plated on medium containing toxic levels of erythromycin and colonies that arose represented bacteria that acquired mutations that constitutively activate the P_*hit*_ promoter (Figure 1C). Thus we named this selection the “*ermC selection*” and the constitutive activating mutations “ON” mutations.

To identify critical residues within HitRS required for signal transduction, *B. anthracis* strains were created in which P_*hit*_ drives expression of two copies of *Escherichia coli relE* (P_*hit*_*relE*) (Figure 1B). The gene *relE* encodes an mRNA endoribonuclease that, when expressed following ‘205-dependent activation of HitRS, cleaves mRNA leading to cell death. In addition, strains were created that harbor one or two copies of *hitRS*, the latter to select for strongly inhibitory variants of HitRS or mutations outside of *hitRS*. Colonies that arose from this selection represent strains containing mutations that render them unable to activate HitRS-dependent signal sensing and gene activation (Figure 1D). The employment of two copies of *relE* excluded mutations within *relE* and enabled preferential isolation of mutations within *hitRS* that inactivate signaling of this TCS. Therefore we named this selection the “*relE selection”* and inactivating mutations “OFF” mutations.

Three types of mutations were isolated from both selections: deletions, frame shifts, and point mutations. Point mutations enabled us to define residues critical for HitRS signal transduction and therefore were the focus of this study. Numerous point mutations were isolated that led to either inactivation or constitutive activation of HitRS, including 40 point mutations that were dispersed in different domains of HitS (2 in the TM domain, 3 in the sensor domain, 17 in the HAMP domain, 9 in the DHp domain, and 9 in the CA domain) (Figure S3A) and 8 point mutations in HitR (3 in the receiver domain and 5 in the DNA-binding domain) (Figure S3B). Among these point mutations, 28 were constitutively activating ON mutations while 20 were inactivating OFF mutations (Figure S3). These point mutations enabled structure-function analysis to interrogate the roles of individual residues and domains and define the molecular basis of HitRS signaling.

### HitS is an intramembrane-sensing HK that detects cell envelope stress

HK sensor domains are highly variable, reflecting the wide variety of input signals that these proteins can sense. Signals perceived by the sensor domain are propagated to the cytoplasm through the TM helices. HitS contains two putative TM helices, the orientation and location of which were consistently predicted by multiple programs including SMART (Letunic & Bork, 2017) and TOPCONS (Tsirigos *et al*, 2015). TM1 (residue 11 to 31) spans from cytosol to exterior while TM2 (residue 47 to 65) spans from exterior to cytosol. These two helices are connected by a 15-amino-acid sensor domain (Figure S3A). Notably, HKs with small sensor domains (≤25 amino acids) have been characterized as intramembrane-sensing HKs (Mascher, 2006, 2014). This group of HKs detect signals within the membrane interface and are often involved in cell envelope stress (Mascher, 2006), which coincides with HitRS being activated by several cell-envelope acting compounds (Mike *et al*., 2014).

Five point mutations from the genetic selections were mapped to this region: two (S25Y and A27E) in TM1 and three (D36V, L42F, and V46G) in the sensor domain. All of these mutations are ON mutations (Figure S3A), indicating that each mutation triggers a sufficient conformational change to enable HitS activation that normally only takes place upon stress detection or ligand binding. Several studies have shown that hydrophilic residues in the TM segment are important for signal recognition (Gushchin *et al*, 2017; Krishnakumar & London, 2007; Zschiedrich *et al*., 2016). Indeed, S25 was substituted by a slightly polar Tyr while the hydrophobic A27 was substituted by a negatively charged Glu, suggesting that it may be a common feature that hydrophilic residues of the TM helices participate in stress detection and signal transduction. There was no clear trend among the three mutations within the sensor domain but all led to constitutive activity of HitS in the absence of any inducers: the polar D36 was mutated to a bulky hydrophobic Val, the bulky hydrophobic L42 was substituted by a relatively less hydrophobic Phe with a larger sidechain, and V46 was mutated to a neutral, small, and flexible Gly (Figure S3A). Very limited structural information is available to dissect the mechanism of ligand recognition and signal detection; however, this short sensor domain likely forms a small extracellular loop and binds ligand directly (Mascher, 2006, 2014). Loop structures can accommodate diverse substitutions, which explains, at least in part, why the drastic changes from these substitutions are tolerated. Nevertheless, these data suggest that these residues identified from the genetic selections are important for ligand binding or stress sensing although the underlying mechanism remains to be elucidated.

### Essentiality of HAMP domain for HitS signaling

The input signal perceived by the sensor domain is subsequently transmitted to the intracellular signaling domains through transducing linkers such as the HAMP domain, which is found in approximately 30% of HKs (Zschiedrich *et al*., 2016). HitS contains a predicted HAMP domain immediately after TM2. To predict the tertiary structure of this domain, homology modeling of this segment (residue 66 to 121) was performed using I-TASSER with default settings (Yang & Zhang, 2015). The closest structural analog was the HAMP domain of *Archaeoglobus fulgidus* Af1503 (PDB ID: 4GN0). The HAMP domain consists of two parallel helices (HAMP1 and HAMP2), which are connected by a flexible loop, and form a homodimeric four-helical parallel bundle (Figure 2).

**Figure 2.**
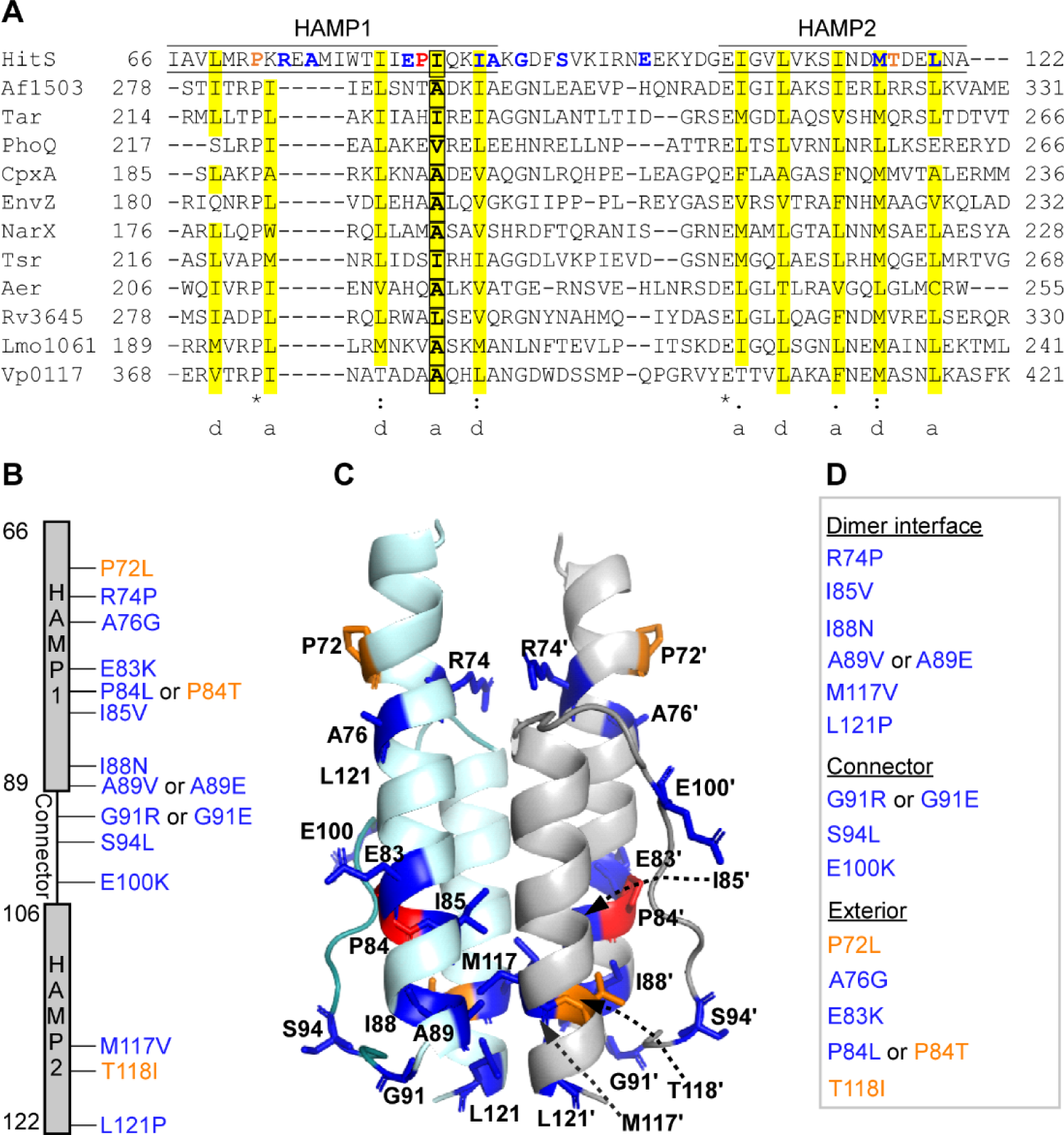
Essentiality of HAMP domain for signal transduction. (A) Multiple HAMP sequences from various histidine kinases, methylated chemotaxis receptors, and adenylyl cyclases were aligned to display the sequence property and conservation pattern of this domain. The two HAMP helices are underlined. The hydrophobic residues at the *a* and *d* positions of the heptad repeat pattern are highlighted in yellow and the residue in HAMP sequence that has been previously reported to be essential for signaling is shown in a box. The residues identified from genetic selections are highlighted in either orange or blue to specify either ON or OFF mutations, respectively. The residue Pro (P84) that can be mutated to either kinase ON or OFF state is highlighted in red. The sequences are the following: HitS, *B. anthracis*; Af1503, *Archaeoglobus fulgidus*; Tar, *Escherichia coli*; PhoQ, *E. coli*; CpxA, *E. coli*; EnvZ, *E. coli*; NarX, *E. coli*; Tsr, *E. coli*; Aer, *E. coli*; Rv3645, *Mycobacterium tuberculosis*; Lmo1061, *Listeria monocytogenes*; Vp0117, *Vibrio parahaemolyticus*. (B) All mutations isolated from genetic selections: OFF mutations are shown in blue while ON mutations are shown in orange. (C) All mutations are mapped onto the homology model of this domain. (D) All mutations were categorized into three groups based on their location in the homodimeric four-helix bundle.

To understand the sequence properties and conservation pattern of this domain, a multiple sequence alignment was performed. HAMP domains are about 50 amino acids long and possess a small number of conserved residues including the Glu residue that marks the beginning of the HAMP2 helix (Figure 2A). Interestingly, HitS contains two sterically restricted Pro residues in HAMP1. The first Pro (P72) is well conserved and its substitution to Leu converted HitS into a constitutively activating ON kinase while the second Pro (P84) is not conserved and its substitution yielded either a kinase-ON (P84T) or kinase-OFF (P84L) state (Figure 2). This signifies the importance of these Pro residues in signal transduction. All HAMP sequences adopt a heptad repeat pattern, in which positions *a* and *d* are occupied predominantly by hydrophobic residues (Figure 2A). These hydrophobic residues are critical for inter- and intramolecular interactions within the four-helix bundle (Hulko *et al*, 2006). Indeed, four hydrophobic residues in this region (I85, I88, M1117, and L121) were identified from genetic selections (Figure 2). I85 is equivalent to a residue that plays a critical role in signal transduction and that a substitution with a larger sidechain at this position promotes signaling while a smaller sidechain substitution compromises autokinase activity in other HKs (Hulko *et al*., 2006). In fact, substitution of I85 to valine with a slightly smaller sidechain completely suppressed the HitS response to stress and inactivated the HitRS system (Figure 2), demonstrating that even small changes at this position can drastically alter autokinase activity. Both M117 and L121 are located in the HAMP2 helix, and it seems likely that substitution of M117 to a bulky Val or L121 to a sterically restricted Pro may disrupt the α-helix conformation resulting in protein instability and loss of autokinase function. Therefore, we conclude that these hydrophobic residues are important for maintaining hydrophobic interactions within the four-helix bundle and are essential for HAMP function.

The flexible connector between the two HAMP helices plays an important role in stabilizing alternating conformations (Airola *et al*, 2010; Ames *et al*, 2008b). It has been shown that a conserved glycine and two hydrophobic residues were the only residues critical for signaling function of a serine chemoreceptor (Ames *et al*, 2008a). Indeed, substitution of the Gly residue (G91) to either positively charged Arg or negatively charged Glu led to complete loss of autokinase function (Figure 2). In addition, substitution of the neutral residue S94 to the hydrophobic bulky Leu or mutation of the acidic E100 to positively charged Lys effectively inactivated the HitRS system (Figure 2), signifying the importance of this connector in TCS signaling through the HAMP domain.

It has been suggested that HAMP domains exist in two conformational states and the transition between the two alternating states is critical for signal transduction (Bhate *et al*., 2015). Therefore, mutations affecting interactions across the close interface within this dynamic four-helix bundle would result in a constitutive ON or OFF conformation, depending on the location of the residues (Swain & Falke, 2007; Swain *et al*, 2009). Consistent with this, we identified seven point mutations located in the dimerization interface: R74P, I85V, I88N, A89V or A89E, M117V, and L121P (Figure 2D), all of which resulted in a kinase-OFF state for HitS (Figure 2B). Collectively, the genetic selections identified 14 critical residues with 17 point mutations within the HAMP domain. The majority of these mutations are OFF mutations (14 out of 17; Figure 2B), highlighting the essentiality of the HAMP domain to HitRS signal transduction.

### All selected point mutants of HitRS have potent growth phenotypes

To obtain a soluble form of HitS membrane protein, the N-terminal 67 amino acids were truncated and the intracellular region of this protein was cloned recombinantly (residue 68 to 352). Homology modeling of the truncated protein was performed using Phyre2 with default settings (Kelley *et al*, 2015). Most crystallization studies of proteins with HAMP domains were performed without this domain, and thus analogous structures only contain part or none of the HAMP domain. The homology model with the highest confidence covers the intracellular domain of HitS from residue 113 to 352. Genetic selections identified 22 point mutations in this region (Figure S3A), 11 of which were selected based on their locations on the structural model for further biochemical characterization: 2 in the HAMP domain, 3 in the DHp domain, and 6 in the CA domain. All 8 point mutations identified within HitR from the genetic selections were spread out in the two domains of HitR (Figure S3B) and subjected to further biochemical characterization to evaluate the effects of these mutation on activities required for signal transduction. Therefore, a total of 21 point mutants including the known phosphorylation-defective variants HitS^H137A^ and HitR^D56N^ (Mike *et al*., 2014) were selected for further study.

First the growth phenotype of these selected suppressor mutants was confirmed. As expected, the parental strain (WT P_*hit*_*ermC*) for *ermC* selection showed no growth in the presence of 20 µg ml^-1^ of erythromycin and reached a similar level of growth after a 15 h lag phase upon ‘205-mediated activation (Figure S4), suggesting that accumulation of *ermC* expression is required for cells to gain resistance against this toxic level of erythromycin. All of the isolated ON mutants showed potent resistance against erythromycin without significant growth delay. Addition of ‘205 provided no evident growth advantage to these mutants under these conditions (Figure S4), suggesting that the erythromycin resistance gene is highly expressed in these constitutively activating ON mutants and inducers are no longer required to turn on the HitRS signaling system.

For the parental strains used in the *relE* selection (P_*hit*_*relE*), ‘205 was added to the medium to activate HitRS, leading to expression of *relE* and disruption of cell growth. Indeed, the *relE* strains grew very poorly with extended lag phases in the presence of ‘205: ∼11 h lag phase for 2x(*relE*) or ∼20 h for 2x(*relE* + *hitRS*) parent strain (Figure S5). By contrast, ‘205 showed no inhibitory effects on any of the OFF mutants isolated and all OFF mutants grew remarkably well regardless of the presence or absence of ‘205 (Figure S5). These results suggest that these OFF mutants were no longer responding to ‘205-mediated activation and their signaling activities were completely abolished.

To confirm the effects of these point mutations on transcription of the *hitPRS* operon, quantitative RT-PCR (qRT-PCR) was carried out to quantify the expression of this operon in these suppressor mutants. As expected, expression of *hitPRS* was upregulated in all ON mutants even in the absence of the inducer ‘205, with the HitS^S141L^ mutation giving rise to the strongest activation for each gene (Figure S6A). Addition of ‘205 activates expression of the *hitPRS* operon in the WT parental strain (WT P_*hit*_*ermC*) but has negligible effects for all ON mutants, which explains why ‘205 provided little growth advantage to these mutants in the presence of erythromycin (Figure S4 and S6B). For some of the OFF mutants, the basal levels of *hitP* were notably lower compared to those in WT parental strains, particularly HitR^Y222D^ and HitR^P155L^ (Figure S6C). Addition of ‘205 induces expression of *hitPRS* in both WT parental *relE* strains, consistent with the poor growth phenotype with extended lag phases observed for these strains (Figure S5 and S6D). As expected, some OFF mutants showed no apparent response to the ‘205 inducer such as HitR^P106S^, HitR^R192C^, HitR^Y222D^, and HitS^M117V^. However, some OFF mutants exhibited a moderate response, particularly the mutants isolated from the *ermC* strain carrying two copies of HitRS (2x(*relE* + *hitRS*)) including HitR^F69S^, HitR^P155L^, and HitS^N248S^ (Figure S6D). All of these latter mutants were isolated from the ectopic copy of *hitRS*, indicating that these OFF mutations are dominant even though the chromosomal copy of *hitRS* was still intact and responsive to inducers in these mutants.

To further confirm the results of the genetic selections, five representative point mutations were reconstructed in *B. anthracis* WT background and the effects of these chromosomal mutations on transcription of the *hitPRS* operon were evaluated using qRT-PCR. Expression of *hitPRS* was constitutively activated in all three ON mutants (HitR^M58I^, HitS^T118I^, and HitS^S141L^) in the absence of the inducer while activation of *hitPRS* was completely abolished in both OFF mutants (HitR^R192C^ and HitS^N248S^) even in the presence of ‘205 (Figure S6E-F). These results demonstrate that the genetic selections are robust and powerful tools to dissect the molecular determinants that are crucial for HitRS signal transduction.

### Critical residues within HitRS stabilize the proteins and facilitate dimerization

Mutations selected in each domain of HitS and HitR were recreated using site-directed mutagenesis and mutant and WT proteins were purified. During the process of protein purification, we noticed some mutant proteins were unstable. Protein misfolding can lead to proteolytic degradation and subsequent protein inactivation. Indeed, four inactive OFF mutants were unstable: M117V and N248S in HitS, and P106S and Y222D in HitR, all of which showed apparent degradation products upon SDS-PAGE. In particular, M117V and Y222D (Figure 3A-B), exhibited susceptibility to proteolysis, indicating that these mutations led to defects in protein folding. Surprisingly, V274A, one of the HitS ON mutants, is partially unstable (Figure 3A). V274 marks the beginning of the D-box and is located in one of the antiparallel β-sheets that hang over the ATP-binding pocket (Figure 3E and S1C). The C-beta branched Val residue is bulky and suitable for β-strand conformation compared to Ala with a small sidechain. Thus this substitution might disrupt the β-strand conformation resulting in defects in protein folding. It is intriguing how this V274A ON mutation promotes phosphorylation at the expense of protein stability.

**Figure 3.**
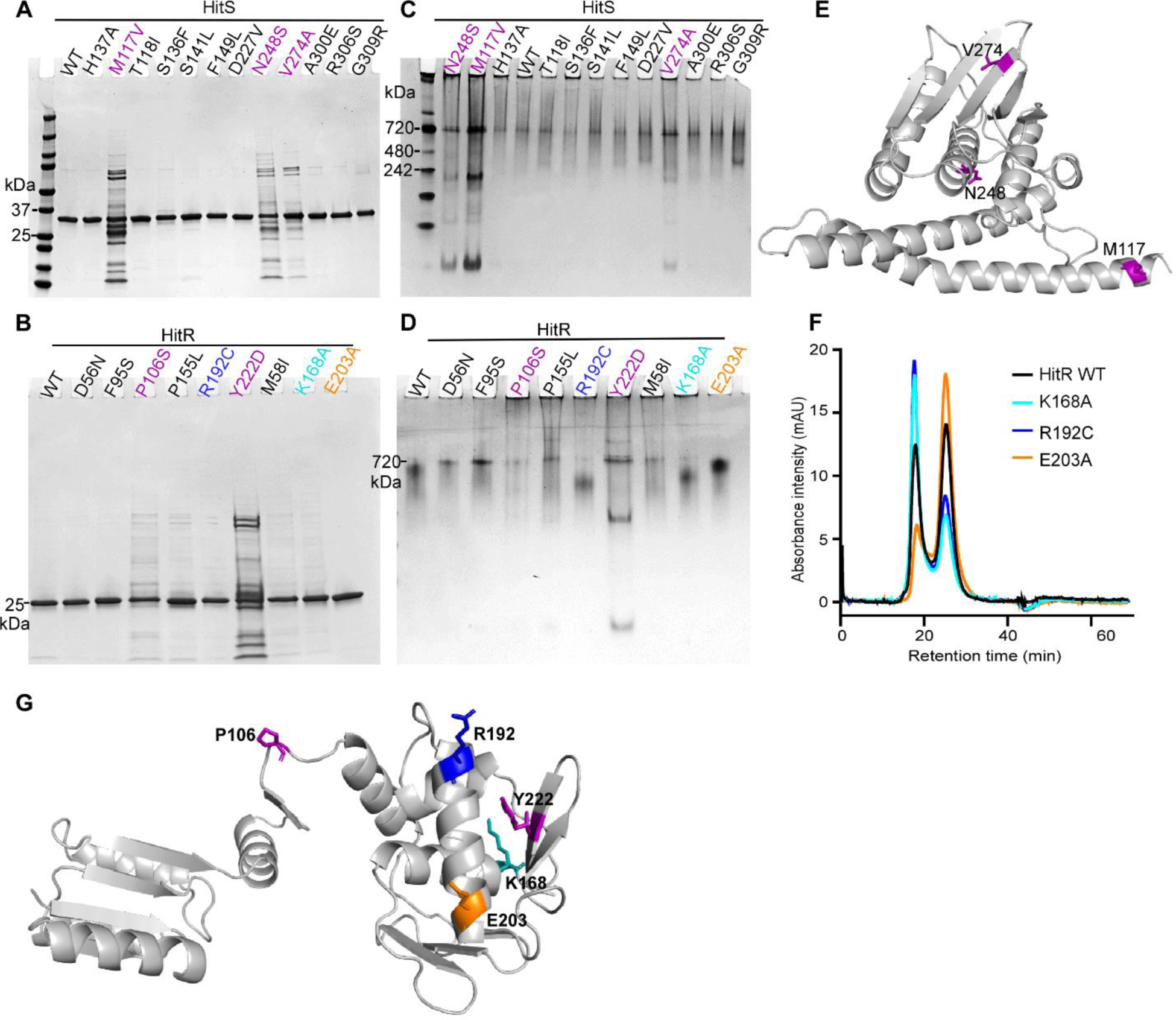
Critical residues within HitRS stabilize protein and facilitate dimerization. To evaluate the effects of HitR and HitS mutations on protein stability and dimerization, WT and mutant proteins were loaded onto SDS-PAGE (A, B) or native gels (C, D). Results shown are WT proteins and all mutants of HitS (A, C) or HitR (B, D) selected for further biochemical characterization. (F) Three HitR mutants affected protein dimerization as determined by size exclusion chromatography. Mutations that affect either protein stability or dimerization are mapped onto the homology models of HitS (E) or HitR (G). HitR model was generated based on *M. tuberculosis* RegX3 (PDB ID: 2OQR), which is in an active dimer form, and only a monomer was shown.

Most HKs and RRs have been demonstrated to function as homodimers (Bhate *et al*., 2015; Zschiedrich *et al*., 2016). The dimer interface is located in the DHp domain of the HK while the receiver domain is dimerized upon RR activation trigged by phosphorylation. To determine the effects of the point mutations on dimerization status, WT and mutant proteins were subjected to non-denaturing native gel electrophoresis to analyze their mobility patterns in the folded state. No significant differences were observed for all HitS mutants compared to WT except that proteolytic degradation of unstable mutants was apparent in the native gel (Figure 3C), which was consistent with the prior results (Figure 3A). All the HitR mutants from the receiver domain formed a relatively sharp band similar to the mutant D56N, which is known to be an inactivating mutant due to loss of phosphotransfer capability (Mike *et al*., 2014). Surprisingly, all four point mutants in the DNA-binding domain of HitR (K168A, R192C, E203A, and Y222D) migrated differently through the gel compared to WT (Figure 3D). Multiple variables could contribute to differences in migration including charge-to-mass ratio, folding status, and physical shape of the protein, which makes it challenging to interpret the different patterns observed. To further examine the dimerization status of these mutants, HitR WT and all of the mutants except the unstable Y222D mutant were subjected to size exclusion chromatography, which separates proteins on the basis of molecular weight. These data showed that the composition of WT HitR was about 45% dimer and 55% monomer in solution (Figure 3F). As expected, the ON mutation K168A promoted dimerization and drove the equilibrium towards the dimeric form with an increase of 25% (Figure 3F), indicating mutation of this polar residue (K168) to a slightly hydrophobic Ala apparently facilitates hydrophobic interactions between monomers. However, it was surprising to note that the OFF mutant R192C promoted dimerization while the ON mutant E203A disrupted dimerization *in vitro* (Figure 3F). This seemed counterintuitive, however some RRs have been shown to form two types of dimers in distinct orientations and only the dimer in the correct orientation is active (Mack *et al*, 2009), which could explain this contradictory observation. In addition, all three substitutions followed the same trend of replacing hydrophilic residues (K168, R192, and E203) with hydrophobic residues (Ala or Cys), indicating that introducing hydrophobic residues at these positions (Figure 3G), particularly K168 and R192, enhances hydrophobic interactions and facilitates dimerization. An important caveat is that RR dimerizes upon phosphorylation and the results may not reflect the dimerization status of these mutants during signal transduction *in vivo*. Thus a thorough evaluation is required to dissect the effects of these mutations on other protein activities such as phosphotransfer and DNA-binding. Nonetheless, these data suggest that residues in the DNA-binding domain interact with the dimer interface of the receiver domain and may affect TCS signaling through modulating HitR dimerization status.

### The autokinase activity of HitS can be modulated in four different manners

To understand how one single-residue mutation alters protein function and locks a protein in a constitutively on or off state, we tested the effects of mutations on different activities intrinsic to the protein including autokinase activity. Consistent with a prior study (Mike *et al*., 2014), substitution of the well-conserved phosphoaccepting histidine residue (H137) to alanine abolishes autokinase activity (Figure 4A-B). Likewise, the two OFF mutations (M117V and N248S) led to a complete loss of autokinase activity (Figure 4A-B). The M117V mutation inactivates autokinase activity potentially by disrupting the α-helix conformation of the HAMP domain while N248S does so likely by disrupting the hydrogen bonds between Asn and ATP adenine resulting in abolished ATP-binding. In addition, two of the ON mutations (D227V and R306S) promoted autokinase activity and several ON mutants showed similar autokinase activity compared to WT (Figure 4A-B). However, we were surprised to note that four ON mutants exhibited significantly reduced autokinase activity: ∼10-fold reduction for S136F, ∼3-fold reduction for F149L, ∼2-fold reduction for V274A, and ∼4-fold reduction for G309R relative to WT. To better understand how these ON mutations affect autokinase activity, autophosphorylation kinetics of HitS WT and ON mutants were monitored for 15 min. Three distinct groupings were revealed: (i) some mutations facilitated the autokinase activity with a higher kinetic rate (A300E, D227V and R306S), (i) some mutations showed minor effects (S141L and T118I), and (iii) some mutations disrupted the autokinase activity with a lower autophosphorylation rate resulting in significantly diminished autokinase yield (S136F, F149L, V274A, and G309R) (Figure 4C-F and table S3). Thus, we conclude HitS autophosphorylation can be modulated in four different manners including abolished activity observed in OFF mutants (Figure 4). Furthermore, these data uncovered four additional residues critical for the autokinase activity: S136/F149 adjacent to the phosphorylation site and V274/G309 from the CA domain, in addition to the well conserved phosphoaccepting His and ATP-binding Asn. However, the observation that ON mutants exhibited reduced autokinase activity appeared contradictory, suggesting that the phosphotransfer rates of these proteins might be altered to compensate for this reduction.

**Figure 4.**
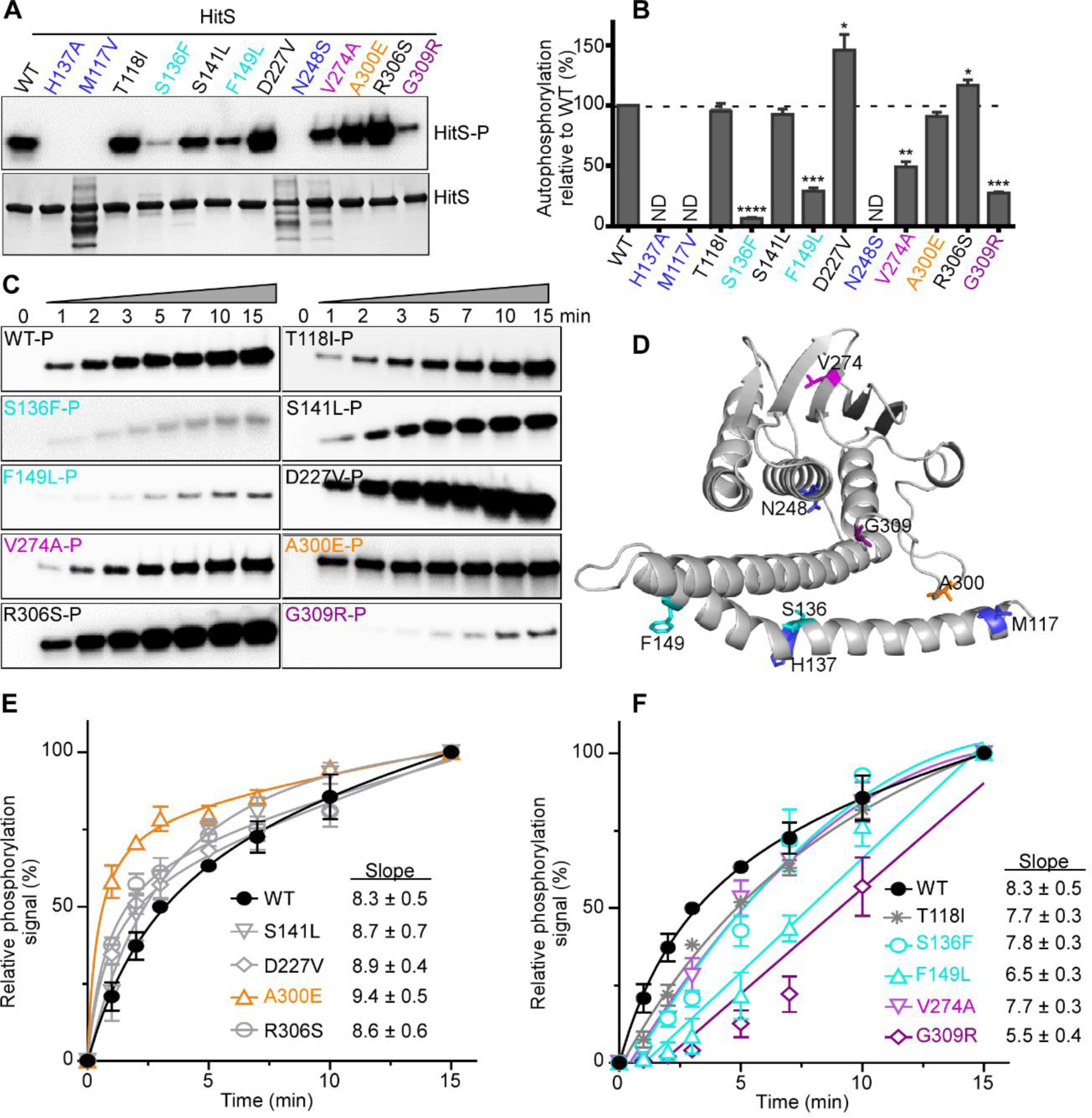
The autokinase activity of HitS can be modulated in four different manners. To evaluate the effects of mutations on HitS kinase activity, the autophosphorylation efficiency of HitS WT and mutants was investigated. (A) Representative phosphor-images (top panel) to show autophosphorylation of HitS WT and mutants that were incubated with ATP [γ-^32^P] for 30 min and quantified using a phosphoimager. The bottom panel is to show the amount of protein used for each reaction in an SDS-PAGE gel. The intensity of the phosphorylation signal was quantified and four independent experiments are shown in (B) (mean ± SEM). Significant differences between WT and each mutant are determined by two-tailed t-test, where \**P* < 0.05, \*\**P* < 0.01, \*\*\**P* < 0.001, and \*\*\*\**P* < 0.0001. (C) Representative phosphor-images to show the kinetics of autophosphorylation by HitS WT or mutants, which was monitored for 15 min. (D) Mutations that affect autophosphorylation are mapped onto a HitS model. (E, F) To better visualize the effects of mutations on HitS autokinase activity, the intensity of the phosphor-signal at different timepoints was quantified and three independent experiments are presented in (E, F) (mean ± SEM). Mutants were organized into two graphs based on their autokinase activity. Data of the first four timepoints (i.e., 0, 1, 2, and 3 min) in E and F were used for slope determination by linear regression analysis.

### Phosphotransfer is the rate-limiting step for signal transduction

Next we examined the impact of all HitS ON mutations on phosphotransfer efficiency. HitS WT or each ON mutant was autophosphorylated with [γ-^32^P]-ATP and phosphotransfer from HitS WT or mutant to HitR WT was then monitored for 15 min. Indeed, all ON mutants transferred the phosphorylation signal significantly faster than WT, including the mutants with defects in autokinase activity such as S136F, V274A, and G309R (Figure 5). This indicates that these mutations are functional trade-offs where the autokinase activity is compensated, at least in part, by faster phosphotransfer. In addition, it is important to note that the phosphotransfer took place in an instantaneous manner. More than 60% of the phosphor signal was transferred from HitS to HitR within 15 seconds (Figure 5A and 5C). Thus we conclude that phosphotransfer from HitS to its cognate regulator HitR is the rate-limiting step for signal transduction.

**Figure 5.**
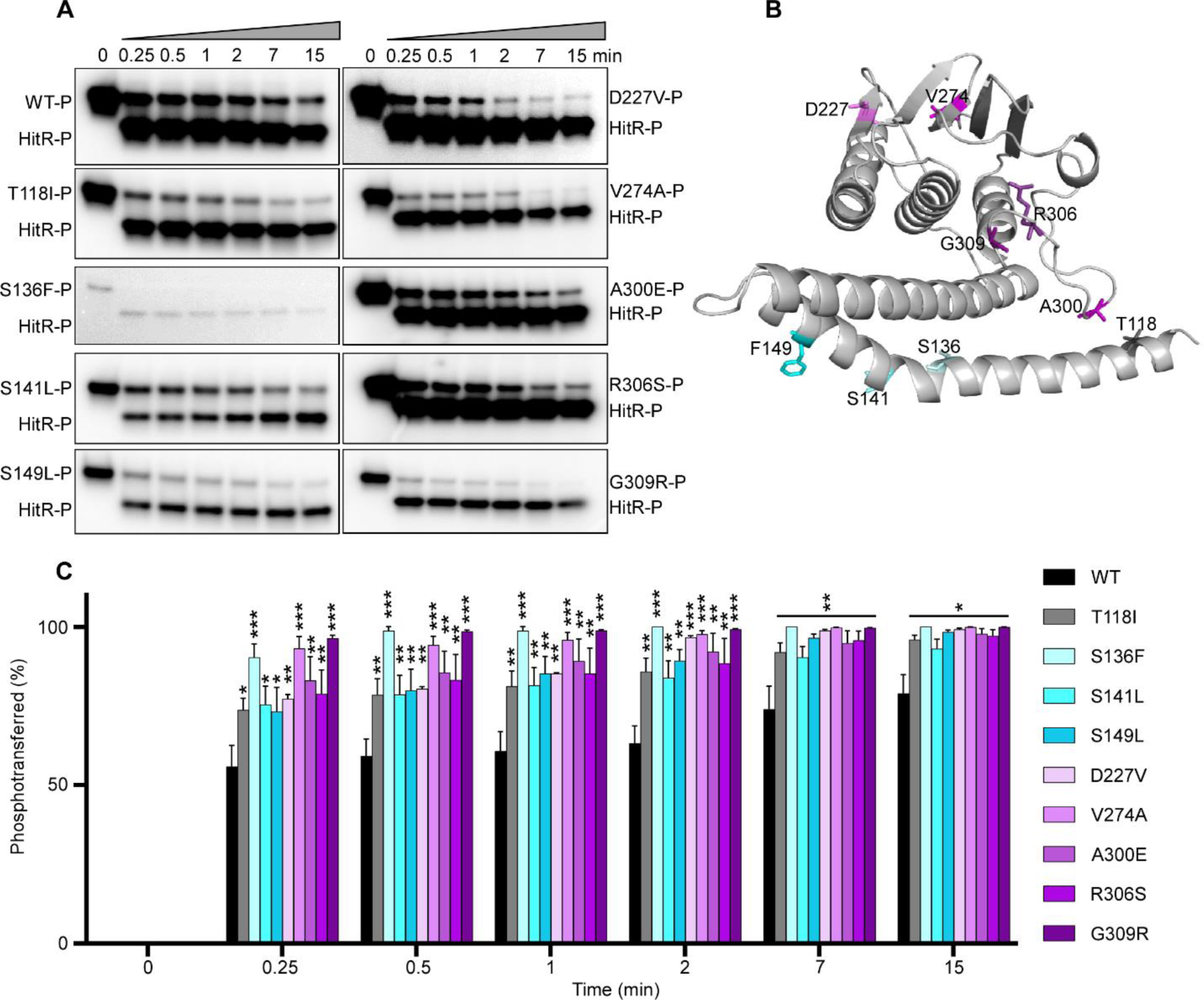
Phosphotransfer is the rate limiting step for signal transduction. To evaluate the effects of HitS mutation on transferring the phosphorylation signal, phosphotransfer efficiency of HitS WT or mutants to HitR WT was examined. (A) Representative phosphor images to show the kinetics of phosphotransfer from HitS WT or constitutively activating mutants to HitR WT, which was monitored for 15 min. (B) Mutations tested are mapped onto a HitS model. (C) The intensity of the lower band (signal transferred) was quantified and relative phospho-signal transferred at different time-points was calculated. Data shown in (C) are from three independent replicates (mean ± SEM). Significant differences determined by two-tailed t-test were observed between WT and each individual activating mutant, where \**P* < 0.05, \*\**P* < 0.01, and \*\*\**P* < 0.001.

### Critical residues required for the phosphatase activity of HitS

Many HKs are bifunctional enzymes that function as both kinases and phosphatases (Batchelor & Goulian, 2003). The autokinase-competent and phosphatase-competent states need to be maintained in balance and modulated in response to specific environmental cues. Importantly, dephosphorylation is not a simple reverse reaction of phosphorylation and may require different residues to achieve this activity. To define the crucial residues required for HitS phosphatase activity, we examined the effects of HitS mutations on dephosphorylation of its cognate regulator HitR. Briefly, the GST-PmrBc fusion protein (Kato & Groisman, 2004) was autophosphorylated and served as a phosphor donor, and the phosphoryl group was subsequently transferred to HitR WT protein. Dephosphorylation of the resultant phosphorylated HitR WT protein was then monitored for 60 min. First, we evaluated the three OFF mutants. H137A abolished the autokinase activity completely (Figure 4A-B); however, the phosphatase activity was intact and comparable to that of WT (Figure 6A and 6C), suggesting that the phosphoaccepting residue is dispensable for the phosphatase activity and HitS is therefore not a reverse phosphatase. The two other OFF mutants (M117V and N248S) showed significantly compromised phosphatase activity (Figure 6A and 6C) with abolished autokinase activity (Figure 4A-B), likely due to instability of these two mutant proteins (Figure 3A). RR dephosphorylation can be catalyzed by either HK-mediated dephosphorylation or auto hydrolysis. The latter is probably why minimal dephosphorylation activity was still observed. Next, we tested the four ON mutants located in the DHp domain, two of which showed drastically diminished activity in dephosphorylation including S141L and F149L (Figure 6A and 6D). We then examined the five ON mutants located in the CA domain, only one (R306S) of which exhibited reduced activity in dephosphorylation (Figure 6A and 6E). When the phosphatase activity of HK is disrupted, this eliminates its ability to remove the phosphoryl group from its cognate RR and the phosphorylated RR can stay active longer thereby promoting signal transduction and gene activation. We conclude these three residues (S141, F149, and R306) are critical for phosphatase activity. Furthermore, these data demonstrated that both HAMP and DHp domains are important for dephosphorylation and not all residues critical for autokinase activity are important for dephosphorylation.

**Figure 6.**
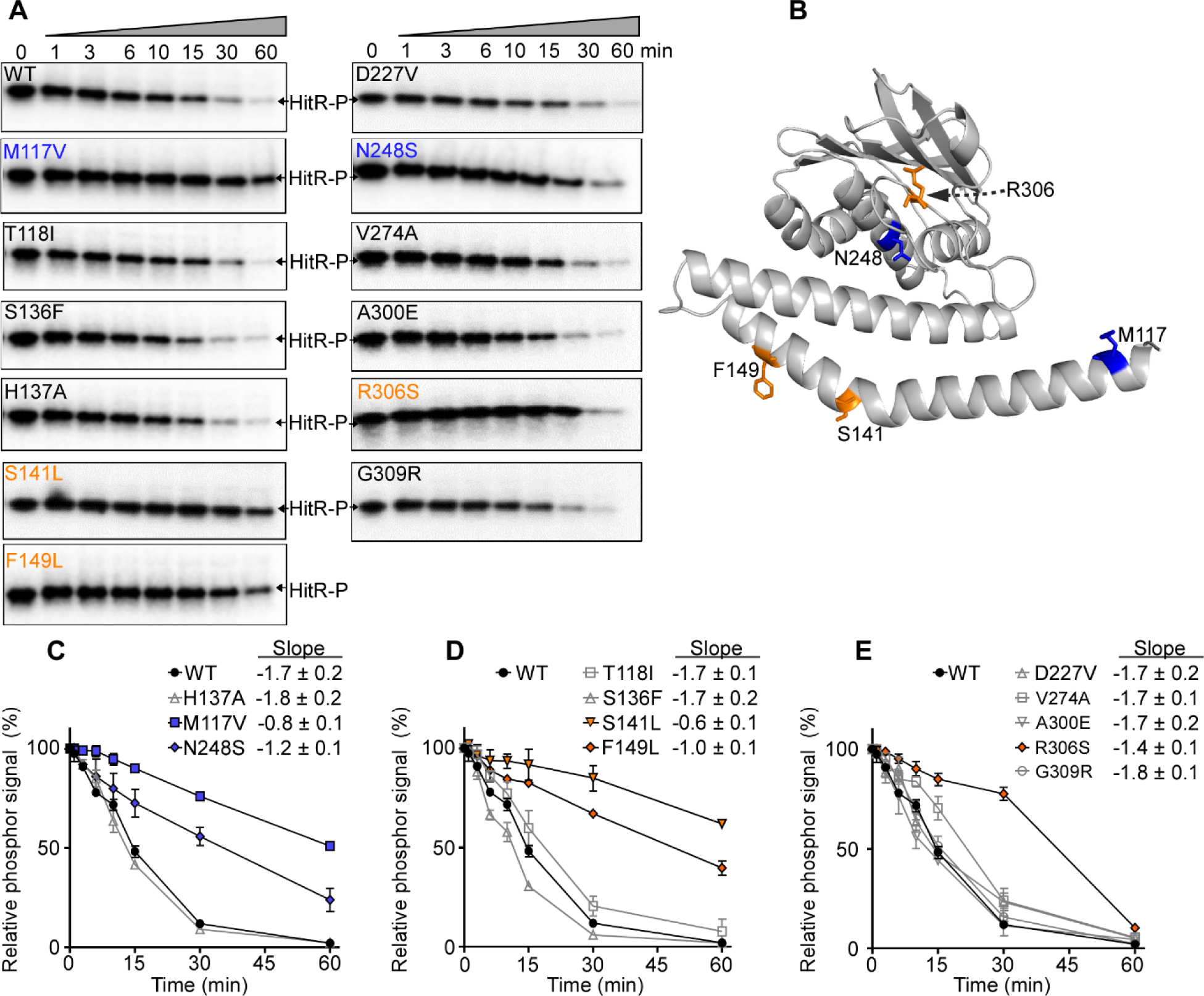
Critical residues required for the phosphatase activity of HitS. To evaluate the effects of HitS mutation on its phosphatase activity, dephosphorylation efficiency of HitS WT and mutants was tested. (A) Representative phospho-images showing the dephosphorylation kinetics of HitS WT or mutants using HitR WT, which was monitored for 60 min. (B) Mutants with altered phosphatase activity were mapped onto the HitS model. (C-E) The intensity of phospho-signal was quantified and relative phospho-signal remaining at different time-points was calculated. Data shown are three independent replicates (mean ± SEM). Mutants were organized into three graphs for optimal visualization. Data of the first four timepoints (i.e., 0, 1, 3, 6 min) for each protein were used for slope determination by linear regression analysis.

### Residues essential for HitR activation and specific interactions within HitR

RRs function as phosphorylation-triggered switches that mediate cellular physiology in response to environmental cues largely through two steps: phosphotransfer from HK to RR and RR-DNA-binding. To examine the effects of HitR mutations on signal reception, the phosphotransfer efficiency was evaluated in HitR WT and HitR mutants. HitS WT was autophosphorylated with ATP [γ-^32^P] and the phosphoryl group was subsequently transferred to HitR WT or mutant proteins. Consistent with a prior study (Mike *et al*., 2014), substitution of the conserved phosphoaccepting residue Asp (D56) to Asn abolishes phosphor signal reception (Figure 7A-B), Two of the three ON mutations (M58I and K168A) showed significantly enhanced activity in phosphotransfer (Figure 7A-B). Among five isolated OFF mutations, only P106S showed ∼50% reduction of activity in phosphotransfer compared to WT (Figure 7A-B), which could be explained by protein instability (Figure 3B). However, it was intriguing that Y222D mutation located in the DNA-binding domain exhibited an equivalent level of phosphotransfer activity compared to WT in spite of being the most unstable HitR mutant (Figure 7A-B and 3B).

**Figure 7.**
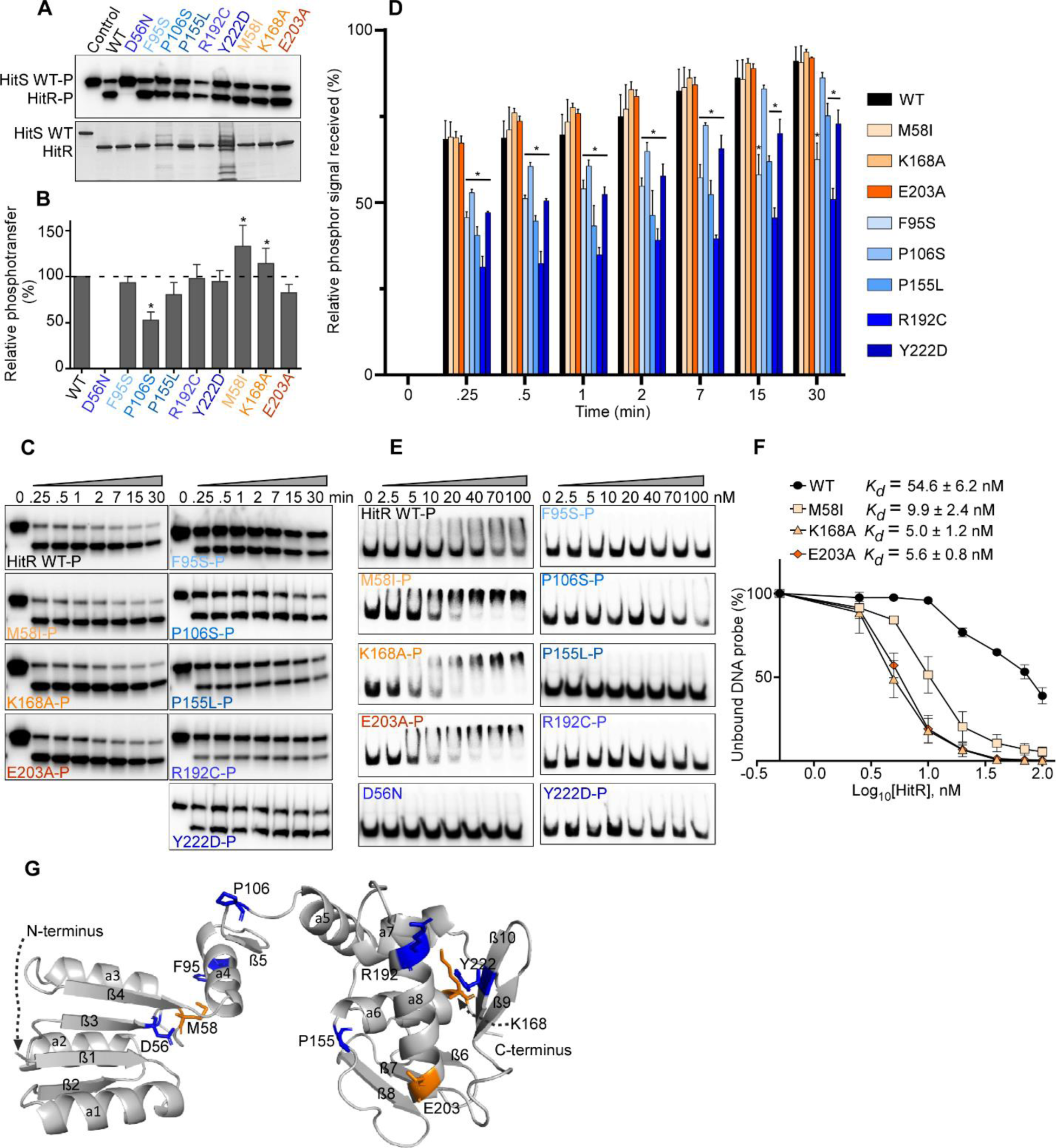
Residues essential for HitR activation and specific interaction within HitR. To examine the effects of HitR mutations on signal reception and DNA-binding, the phosphotransfer efficiency and DNA-binding affinity of HitR WT or mutants were tested. (A, B) Phosphotransfer efficiency from HitS WT to HitR WT or mutants was quantified. (A) The top is a representative phosphor image and the bottom is an SDS-PAGE gel showing the amount of protein used for each reaction. The intensity of radioactive signal was quantified and averages from four independent experiments are shown in (B) (mean ± SEM). Statistical significance was determined by two-tailed *t*-test, where \**P* < 0.05. (C) Kinetics of phosphotransfer from HitS WT to HitR WT or mutants was monitored for 30 min. Representative images are shown. The intensity of the lower band (phosphotransferred) at each timepoint was quantified. Presented are averages from three independent replicates (D) (mean ± SEM). (E) Representative images to show DNA-binding of HitR WT or mutants to its target promoter evaluated by electrophoretic mobility shift assay (EMSA). (F) The band intensity of unshifted DNA probe (lower band) was quantified using GelQuantNET. All data points from three independent experiments were plotted and subjected to *K*_d_ determination using GraphPad Prism 8 (mean ± SEM). (G) All point mutants tested are mapped onto a HitR structure model. All blue colors indicate OFF mutation while orange colors indicate ON mutation.

To better understand how these mutations affect phosphor signal reception, the kinetics of phosphotransfer from HitS WT to HitR WT or mutants was monitored for 30 min. None of the ON mutants exhibited an accelerated rate of phosphotransfer while all OFF mutants displayed significantly slower kinetic rates relative to WT HitR (Figure 7C-D). M58 is located in the loop between β3 and α3 of the receiver domain (Figure 7G), and loop structures can tolerate a large number of diverse substitutions, which is probably why substitution of M58 to a bulky Ile showed no evident effects on protein stability (Figure 3B). In addition, M58 is in very close proximity to the phosphoaccepting residue D56 (Figure 7G) and the substitution to Ile appeared to facilitate phosphotransfer, indicating that M58 is involved in phosphorylation signal reception. K168 marks the C-terminal end of α6 in the DNA-binding domain and its substitution to Ala had no notable effects on protein stability (Figure 3B) but promoted phosphotransfer (Figure 7A-B), suggesting that the DNA-binding domain interacts with the receiver domain and facilitates phosphorylation reception. These data also indicate that α6 helix may be part of the interface of these two domains and important for their interaction between these two domains. P155 marks the end of the loop between β8 and α6 in the DNA-binding domain (Figure 7G), and its substitution to Ser was permitted in the loop structure, resulting in a mutant protein with comparable stability to WT (Figure 3B). However, the tight turn created by Pro was likely destroyed by Ser substitution resulting in defects in phosphotransfer (Figure 7A-D). This indicates that this flexible loop not only enables conformational changes transmitted from the receiver domain to the effector DNA-binding domain, but also facilitates phosphotransfer from HK to RR. Both R192 and Y222 are located in the DNA-binding domain. R192 resides in helix α8 and Y222 is in the last β-strand. Interestingly, mutation of either residue affected reception of the phosphoryl group, exemplifying the importance of the DNA-binding domain during phosphotransfer and interaction between the two domains of HitR.

Next, we investigated the effects of HitR mutations on HitR-DNA-binding ability using an electrophoretic mobility shift assay (EMSA). HitR WT or mutant protein was first activated by autophosphorylation, and the binding affinity to the target promoter was then evaluated. HitR WT binds DNA with an affinity of ∼55 nM, and as expected, the D56N mutation led to a complete loss in DNA-binding. Remarkably, no visible band-shift was observed for all five OFF mutations with up to 100 nM of protein tested except P106S, which showed a smear with the highest protein concentrations (70-100 nM), indicative of a protein-DNA complex (Figure 7E). Most of the OFF mutants showed a relatively slower rate in phosphorylation reception without much difference in the final outcome compared to WT, with the only exception of P106S that showed a 50% reduction in phosphotransfer within 30 min (Figure 7A-D). However, the effects of these mutations on protein-DNA-binding were strikingly disruptive (Figure 7E), further confirming that even a small difference in the kinetic rate can lead to severe defects in protein function, consistent with phosphotransfer being the rate limiting step for signal transduction. As anticipated, all ON mutants exhibited much higher affinity to the target promoter. M58I mutation facilitates phosphotransfer from HitS to HitR, which in turn promotes DNA-binding with a 5-fold increase in binding affinity. Both ON mutations (K168A and E203A) from the DNA-binding domain had no significant effects on phosphotransfer kinetics. However, these two mutations dramatically enhanced protein-DNA-binding with 10-fold higher affinity compared to that of WT (Figure 7E-G). We hypothesized that these two ON mutants might bypass phosphorylation and bind DNA with greater affinity without phosphorylation-mediated activation.

To test the essentiality of HitR activation through phosphorylation for HitR-DNA-binding, we repeated EMSA experiments using WT HitR and the three ON mutants in the absence of phosphorylation. HitR WT or mutant were incubated directly with radioactively labelled DNA probe and the binding affinity was examined. Surprisingly, HitR WT binds to DNA with comparable affinity regardless of phosphorylation activation (Figure S7), likely due to overexpression of HitR in *E. coli* that led to a conformational transition from an inactive to active-like state as observed for KdpE previously (Narayanan *et al*, 2014). However, in the absence of phosphorylation, the DNA-binding activity of the M58I mutant was completely abolished with up to 800 nM of the mutant protein tested. K168A showed similar DNA-binding activity as WT while E203A only preserved minimal activity (Figure S7), which could be explained by the influence of these mutations on HitR dimerization status: K168A facilitated dimerization while E203A disrupted dimerization *in vitro* (Figure 3F), signifying the importance of dimerization in HitR activation. Furthermore, it is clear that phosphorylation-trigged activation is crucial for HitR-DNA-binding. Taken together, we conclude that the receiver and DNA-binding domains communicate through critical residues as they work together to ensure HitR phosphorylation and downstream gene activation in response to specific stimuli.

### Residues critical for HitS-HitR interaction

It is noteworthy that the HitS ON mutation of F149 to Leu in the RR-binding interface led to potent activation of HitRS signaling (Figures S1A, S4E, S6A and Table S3). This is a typical functional trade-off mutation that affects all three activities with diminished autophosphorylation, enhanced phosphotransfer, and disrupted dephosphorylation (Table S3), underscoring the significance of the RR binding interface in all three catalytic reactions of HitS. We reasoned that this mutant, along with some other mutants from Helix 1, would affect HitS-HitR interaction. To test this idea, we determined HitS-HitR binding affinity *in vitro* using microscale thermophoresis. The two WT proteins bound to each other with an affinity of 385nM (Figure S8A), and activated HitS modestly enhanced the binding affinity with a *K*_*d*_ value of 270 nM (Figure S8B). The difference was not dramatic but could be physiologically relevant during HitRS signaling *in vivo* since some RRs even exhibit a reduced affinity for their cognate kinases upon phosphorylation (Li *et al*, 1995). Surprisingly, both HitR^S141L^ and HitR^F149L^ drastically disrupted the protein-protein interaction. Specifically, when the mutants were not activated, the binding affinity was 4-6 times lower compared to WT (Figure S8C and S8E). When HK is not activated through autophosphorylation, it is in phosphatase-competent state. These data explained why the phosphatase activity of these two mutants (HitR^S141L^ and HitR^F149L^) was disrupted (Figure 6A and 6D). By contrast, activation of these two mutants mediated by autophosphorylation improved their binding affinity to HitR, although still significantly weaker than WT (Figure S8D and S8F). These data indicate that these two residues (S141 and F149) of HitS are important for HK-RR interaction particularly during dephosphorylation.

## Discussion

Recent structural and biochemical work has provided valuable signaling models and substantially deepened our understanding on the molecular basis of signal transduction from external input domains to cytoplasmic output domain. Heretofore, there are more than 600 three-dimensional structures of HKs and RRs available; however, most of these structures are for individual domains, particularly for the membrane-bound HKs. It is challenging to obtain high-resolution structures of full-length proteins due to solubility, flexibility, and dynamics of the sensor HKs. Individual domains have inherent features and functional modes, but their interactions with other partners are crucial for specific signaling pathways and regulatory mechanisms. In this study, robust and unbiased genetic selections enabled selection of point mutations within HitRS that constitutively switch on or off signal transduction of this TCS and these point mutations were further characterized systematically. Our data demonstrated the effects of these mutations on diverse activities intrinsic to TCS signaling, defined the critical residues that are involved in HitRS signal transduction, determined phosphotransfer as the rate-limiting step for signal transduction, and shed light on the signaling mechanism of each individual domain and the TCS as a whole.

### Hydrophobic interactions within HAMP domain

HAMP domains in HKs are typically located immediately after the C-terminal transmembrane helix and function as signal transducing modules that couple conformational changes of sensory domains to the catalytic activity of the kinase core domains (Bhate *et al*., 2015; Zschiedrich *et al*., 2016). HAMP domains can be swapped among proteins without compromised functionality (Appleman *et al*, 2003; Zhu & Inouye, 2003), indicative of a conserved signaling mechanism. A few models have been proposed for the mechanism of signal transduction through the HAMP domain (Airola *et al*., 2010; Hulko *et al*., 2006; Parkinson, 2010; Stewart, 2014; Swain & Falke, 2007; Swain *et al*., 2009). These models differ in many aspects but share one commonality: the HAMP domain shuttles between two distinct conformations, which represent two opposing signaling states and are stabilized by different subsets of conserved residues (Zschiedrich *et al*., 2016). Indeed, we isolated a total of 17 point mutations from genetic selections, either constitutively activating ON (3) or inactivating OFF mutations (14), within this 50-residue HAMP domain (Figure 2). Each individual point mutation induces conformational changes sufficient for switching the signaling function to either an on or off state, which exemplifies the dynamics, flexibility, and interchangeable nature of the HAMP domain. Five hydrophobic residues identified are located at the dimer interface (I85, I88, A89, M117, and L121) and any mutations to neutral, hydrophilic, or even slightly less hydrophobic residues would drastically affect conformation of this domain resulting in loss of function (Figure 2). Thus, it is clear that these hydrophobic residues pack together in the interior of the helix bundle and stabilize protein conformation through hydrophobic interactions. Furthermore, among three constitutively ON mutations isolated from the HAMP domain, two were substitutions from neutral residues (P72 and T118) to hydrophobic residues (Leu and Ile, respectively; Figure 2). Both Leu and Ile are highly hydrophobic and custom-made for introducing additional hydrophobic effects to stabilize the kinase-competent conformation of this helix bundle. In conclusion, this and other studies demonstrated the importance of hydrophobic residues in HAMP function and the essentiality of this domain in TCS signaling.

### The kinase core: DHp and CA domains and their interaction

The DHp domain forms a homodimeric antiparallel four-helix bundle with two α-helices joined by a hairpin loop (Figure S1A-B). HitS possesses a HisKA subfamily DHp domain. Three catalytic reactions take place at this domain: (i) histidine autophosphorylation, (ii) phosphotransfer from HK to its cognate RR, and (iii) dephosphorylation of the phosphorylated RR. This helix bundle can be divided into three segments based on prior DHp sequence analysis (Bhate *et al*., 2015) (Figure S1B). Below we summarize the role for each segment and the effects of mutations in that segment have on the function.

First, the top region serves as the binding site for the Gripper fingers of CA during autophosphorylation. It switches between symmetric and asymmetric conformations, which correlate with phosphatase-competent and kinase-competent states, respectively (Bhate *et al*., 2015). Five ON mutations were isolated from this region (S136L, L184F, L185R, T188I, and L189P) (Figure S1B). The sidechain of S136 or T188 likely forms a hydrogen bond with the protein backbone and its substitution to a hydrophobic residue (Leu or Ile) introduces hydrophobic interactions within the four-helix bundle and enables HAMP helices to shift conformation towards a kinase-competent state. On the other hand, the hydrophobic Leu (L184, L185, or L189) was mutated to a relatively less hydrophobic (L184F), hydrophilic (L185R), or sterically restricted residue (L189P) (Figure S1B), all of which resulted in asymmetric kinase-on conformation.

Second, the highly symmetric central core follows immediately after the well-conserved phosphoaccepting His and functions as the docking site for the cognate RR during either phosphotransfer or dephosphorylation (Bhate *et al*., 2015). The conserved (T/N)-P dipeptide is known to form a kink for helix bending that allows the N-terminus of the HAMP1 helix to adopt multiple conformations during signaling (Bhate *et al*., 2015). HitS contains a Ser, the preferred substitution residue for Thr, along with a conserved Pro at this position. Either residue could be mutated (S141L or P142S) to disrupt this tight turn on the protein surface and lock the kinase in a constitutively on conformation (Figure S1B). In addition, L177 located in the interior of the four-helix bundle was mutated to a much less hydrophobic Ala and this mutation also triggered sufficient conformation changes from stable symmetric to asymmetric state, indicating the involvement of L177 in the dimer interface. Furthermore, both S141L and F149L mutations disrupted the interaction of HitS with its cognate regulator HitR particularly during dephosphorylation (Figures S8C and S8E) resulting in potent activation of HitRS signaling (Figure S4D-E). These data illustrate the importance of this segment for HK-RR interactions and TCS signal transduction.

Third, the bottom part of the bundle is the continuity of the RR-binding interface joined by a hairpin loop. The sequence and length of this region are highly variable (Figure S1A), reflecting sequence-specific interactions for recognition of the cognate RR. In conclusion, these results strongly support prior DHp sequence analysis (Mechaly *et al*, 2014) and demonstrate that the symmetry-asymmetry transition is a key feature of signal transduction through the DHp domain.

The CA domain is well conserved with N, G1 (or D), F, G2, G3 sequence motifs (Figure S1A and S1C), which are all defined by the critical residues within these boxes and are all involved in ATP-binding and catalysis. In between the F and G2 boxes, a flexible loop called the ATP-lid covers the ATP-binding site (Figure S1A and S1C). The flexibility of the ATP-lid is important for ATP-binding as well as interacting with the DHp domain, which allows the CA domain to bind to different regions of DHp during multiple catalytic reactions depending on DHp conformation and the catalytic status of the CA domain (Albanesi *et al*, 2009; Marina *et al*, 2001; Marina *et al*, 2005; Trajtenberg *et al*, 2010). A Gripper helix with four hydrophobic residues named Gripper fingers, located within the G2-box (Figure S1A and S1C), was recently defined and works together with a Phe in the F-box to mediate the interaction with the DHp domain (Bhate *et al*., 2015). Interestingly, most of the point mutants identified within the CA domain were located in this region: four in the ATP-lid (A300E, N305K, R306S, and G309R) and two within the Gripper helix (A316E and K320E) (Figure S1A and S1C). Each individual mutation triggers structural changes of the CA domain and the interacting partners of CA and shifts the conformation equilibrium of the entire HK towards kinase-on state, highlighting the flexibility and versatility of these two motifs. Three of these mutants were characterized *in vitro*: both A300E and R306S promoted autokinase and phosphotransfer activities while G309R enhanced phosphotransfer with drastically diminished autophosphorylation. In addition, both A300E and G309R had no impact on phosphatase activity while R306S significantly disrupted dephosphorylation of the phosphorylated HitR, demonstrating the involvement of CA domain in all three catalytic activities. In conclusion, these data along with other structural analyses (Bhate *et al*., 2015; Jacob-Dubuisson *et al*., 2018; Zschiedrich *et al*., 2016) support a model in which the CA domain needs to adopt several positions relative to the DHp domain during different catalytic states of the HK, and the interaction between DHp and CA domains mediated by the Gripper helix are critical for TCS signaling.

### The receiver and DNA-binding domains of RR and regulation mechanism

The receiver domains share a conserved (βα)5 fold (Figure S2A and S2C) and function as phosphorylation-dependent switches to modulate the activity of the effector domain using distinct inter- and/or intramolecular interactions in the inactive and active states (Gao *et al*, 2007). Some RRs have been reported to dimerize in two different orientations: one involves the α4-β5-α5 surface and the other involves the α1/α5 surface and only the α4-β5-α5 dimer is functional upon activation (Mack *et al*., 2009). The majority of the exposed sidechains on the α4-β5-α5 surface are hydrophilic, suggesting that any interactions through this interface are likely to be dynamic (Barbieri *et al*, 2010; Mack *et al*., 2009). Indeed, two residues (F95 and P106) identified from the *relE* selection are located in this region: both mutants (F95S and P106S) were still capable of accepting phosphor signals from HitS with rather slower kinetic rates but the DNA-binding ability of both mutants was nearly abolished (Figure 7 and Table S3), indicating the dimerization step between phosphotransfer and DNA-binding is likely disrupted. The hydroxyl group in the sidechain of Ser is fairly reactive and may form hydrogen bonds with the polar residues in the α4/α5 helices and disrupt the dynamics and flexibility of this α4-β5-α5 dimer interface (Figure S2C), which in turn prevents HitR from dimerizing through this interface thereby abrogating HitR activation (Figure S4-5). These data suggest the dynamics and flexibility of this α4-β5-α5 interface may be important for RR activation through dimerization in the correct orientation. A conserved Met (M58) at two residues after β3 strand (Figure S2A and S2C) was identified from the *ermC* selection and its mutation to a bulky and mostly hydrophobic Ile promoted the phosphoryl group transfer from His to Asp (Figure 7). It is unclear how Ile substitution at this position facilitates phosphotransfer. There are two possibilities: (i) introducing hydrophobic residues to protein surfaces can stabilize a protein through improved water–protein interactions as reported recently (Islam *et al*, 2019), or (ii) Ile is in close proximity to the dimer interface and can facilitate dimerization through hydrophobic effects. Nonetheless, M58I substitution led to conformational changes of the receiver domain resulting in activation of the associated DNA-binding domain and downstream gene transcription (Figure S4K and S6).

RRs with DNA-binding domains can be categorized into four subfamilies: OmpR/PhoB, NarL/FixJ, NtrC/DctD, and LytR/AgrA. HitR belongs to the largest OmpR/PhoB subfamily with a conserved winged helix-turn-helix fold (Figure S2D). Helix α8 and the last two β-strands (β9/β10) are critical for DNA-binding: α8 helix recognizes the specific DNA sequence and inserts into the major groove of DNA while the β-hairpin (β9/β10) binds in the minor DNA groove (Blanco *et al*, 2002). Two residues were identified within the α8 helix by the genetic selections: R192 and E203 and one residue within strand β10: Y222. The sidechains of both polar residues (R192 and E203) likely participate in hydrogen bonding with specific bases in the major DNA groove. Interestingly, substitution of R192 to a hydrophobic Cys led to complete loss of DNA-binding while mutation of E203 to a hydrophobic Ala enhanced DNA-binding with a 10-fold increase in binding affinity compared to WT (Figure 7). Hydrophobic residues (Cys and Ala) can also interact with DNA bases and stabilize protein-DNA complexes through hydrophobic interactions although the completely opposite effects of these two mutations seemed counterintuitive. However, R192C disrupted phosphotransfer while E203A showed no evident effects on phosphotransfer, indicating that R192 but not E203 in helix α8 of the DNA-binding domain may interact with the receiver domain and facilitate transfer of the phosphoryl group from HK to RR. R192C mutation disrupted this interaction leading to diminished phosphotransfer and abolished DNA-binding while E203A enhanced DNA-binding likely through hydrophobic effects. Y222 is located only six residues away from the C-terminus of HitR, however, its substitution to acidic Asp drastically disrupted the β-strand conformation and led to protein instability and complete loss of DNA-binding (Figure 3B and 7E, Table S3). This mutant was still able to receive phosphoryl group from HK with a lower kinetic rate (Figure 7A-D), indicating that the β-hairpin may be not involved in phosphotransfer but might play a critical role in DNA-binding.

Collectively, this study provides a detailed characterization and structure-function analysis of an entire TCS, defines molecular determinants of each domain for both HK and RR, reveals residues critical for various activities intrinsic to TCSs, uncovers interaction specificity among different domains and between the HK and RR, and extends our understanding of the molecular basis of signal transduction through TCSs. In addition, all constitutively ON point mutants identified within these two well-conserved signaling proteins could be useful for studying fundamental mechanisms of signaling in other TCSs with unknown targets or unknown partners. These ON mutations could also serve as a blueprint for developing biotechnology tools suitable for synthetic biology engineering that connects sensory modules with signaling outputs (Ninfa, 2010). Given that antibiotic resistance is one of the most significant threats to global health, novel antimicrobial therapeutics are in desperate need. It is reasonable to think that the well conserved HAMP domain found in many sensor and chemotaxis proteins could be the top target for rational inhibitor design to disrupt the hydrophobic interactions within the four-helix bundle and disrupt the signal transmission that is required for many biological processes. This study along with other findings pave the way for developing novel antimicrobials or adjunctive treatments that target signal transduction in infectious pathogens.

## Materials and Methods

Materials and methods are described in the SI Appendix, including growth conditions, DNA manipulation and strain construction, genetic selections, growth curves, qRT-PCR, homology modeling, protein expression and purification, size exclusion chromatography, autophosphorylation assay, phosphotransfer assay, dephosphorylation assay, EMSA, and microscale thermophoresis assay.

## Acknowledgments

We thank Jocelyn Simpson for her help on *ermC* selection. We thank Dr. Heather Kroh for her technical advice and Dr. Maria Hadjifrangiskou for sharing the GST-PmrBc construct. We thank members of the Skaar Laboratory for critical comments of the manuscript. M.L.C, S.J.I, C.J.L, A.M.T, S.M.C, G.H.H, H.K.L, J.D.R and D.L.S. were supported by the Grove City College Swezey Fund and the Jewell, Moore, and MacKenzie Fund. Work in the Skaar Lab was funded by National Institutes of Health grant R01 AI73843 to E.P.S and H.P. was supported by T32 HL094296.

## Author Contributions

H.P. and E.P.S. conceived and designed the experiments, H.P. acquired all experimental data exclusive of *B. anthracis* strain construction and genetic selections. M.L.C, S.J.I, C.J.L, A.M.T, S.M.C, G.H.H, H.K.L, J.D.R and D.L.S. constructed *B. anthracis* strains used for genetic selections and the chromosomal *hitRS* point-mutants. M.L.C, C.J.L, and D.L.S. conducted genetic selections. H.P. drafted the manuscript, HP, D.L.S, and E.P.S edited the manuscript, and all authors reviewed the manuscript.

